# Tristetraprolin promotes survival of mammary progenitor cells by restraining TNFα levels

**DOI:** 10.1101/2023.04.04.532205

**Authors:** Stedile Micaela, Lara Montero Angela, García Solá Martín Emilio, Goddio María Victoria, Beckerman Inés, Bogni Emilia, Ayre Marina, Naguila Zaira, Coso Omar, Edith C. Kordon

**Affiliations:** Instituto de Fisiología, Biología Molecular y Neurociencias (IFIBYNE), CONICET-UBA, Buenos Aires, 1428, Argentina; Bioterio Central, Facultad de Ciencias Exactas y Naturales (FCEN), Universidad de Buenos Aires (UBA), Buenos Aires, 1428, Argentina; Departamento de Fisiología, Biología Molecular y Celular (DFBMC), Facultad de Ciencias Exactas y Naturales (FCEN), Universidad de Buenos Aires (UBA), 1428, Argentina; Departamento de Química Biológica (DQB), Facultad de Ciencias Exactas y Naturales (FCEN), Universidad de Buenos Aires (UBA), 1428, Argentina

## Abstract

Tristetraprolin (TTP) is a RNA binding protein that destabilizes mRNA of factors that up-regulate proliferation, invasiveness and inflammation. Here we show that TTP expression is higher in mammary progenitor cells than in other cell populations, and that reducing its levels impairs mammary gland morphogenesis *in vivo* and mammosphere formation in culture. Knocking down TTP in stem-like HC11 mouse mammary cell line increased inflammatory cytokine mRNAs and signaling cascades involving NFκB, STAT3 and MAPK p38 activation, which led to apoptosis. Importantly, TNFα overexpression and the consequent p38 phosphorylation would be the leading cause of progenitor cell death upon TTP expression restriction. Taken together, our results reveal the relevance of negative posttranscriptional regulation on TNFα, exerted by TTP, for the maintenance of the progenitor cell compartment in the mammary gland.

## INTRODUCTION

The mammary epithelium entails two main cellular lineages: luminal cells that border the ductal and alveolar lumen, and myoepithelial or basal cells, which are located next to the basement membrane (Watson and Khaled, 2008). The proliferative changes within this epithelium are fueled by embryonic pluripotent mammary stem cells (MaSCs), capable of originating both basal and luminal lineages, and by unipotent progenitor cells (*i.e.* luminal and basal) that amplify and maintain each of these compartments in the post-natal gland throughout the successive reproductive cycles of the adult female mouse (Visvader and Stingl, 2014). Starting at puberty, a tree of branching ducts driven by the terminal end buds (TEBs) located at their ends, advance until they reach the limits of the mammary fat pad, when these structures disappear (Williams and Daniel, 1983). In each pregnancy, progenitor cells initiate the secretory lobuloalveolar compartment, which is the main source of milk production during lactation. After weaning, these differentiated structures regress due to massive cell death and tissue remodeling. This involution process is associated with inflammatory pathways that involve activation of transcription factors as STAT3 and NFkB (Inman et al., 2015; Watson and Kreuzaler, 2011). However, a pregnancy-induced mammary epithelial cell (PI-MEC) population that originates in the first pregnancy and lactation cycle do not die out during involution and persists throughout the rest of the female mouse life. These cells possess stem cell-like features, are at least partly responsible for subsequent lobular development and may behave as cancer stem cells (Wagner and Smith, 2005).

Tristetraprolin (TTP) is a protein encoded by *Zfp36* gene that binds to sequences enriched in Adenine and Uracile (AU-binding protein, AUBP) in the 3′unstranslated regions (3’UTR) of specific mRNAs, leading mostly to their decay. However, it has been reported that TTP is also able to modulate expression of different targets also at transcriptional and translational levels (Guo et al., 2017; Kovarik et al., 2021). TTP controls many inflammation-associated mRNAs and is required for the resolution of inflammation as well as immune homeostasis However, its activity is very dependent on the cellular context (Kovarik et al., 2021). In addition, TTP has been described as a tumor suppressor, because it negatively regulates the expression of multiple genes involved in cell proliferation and cycling, cell death inhibition, angiogenesis induction, epithelial to mesenchymal transition and invasiveness (Guo et al., 2017).

We have previously reported that TTP expression is induced by prolactin in differentiated mammary epithelial cells (Goddio et al., 2012) and its presence is required for lactation maintenance in female mice, since lactating TTP^fl/fl^ x Wap-Cre bitransgenic females (MG-TTP KO) showed premature involution (Goddio et al., 2018). Here, we show that TTP down-regulation affects not only the fully differentiated mouse mammary epithelium, but also the progenitor cell survival resulted compromised both *in vivo* and in culture. We found that this deleterious effect would be mostly due to the dramatic TNFα overexpression and activation of a p38 dependent pro-apoptotic pathway.

## RESULTS

### TTP/*Zfp36* down-regulation induces mammary progenitor cell impairment ***in vivo***

As we have previously observed during the first lactation period of MG TTP-KO females (Goddio et al., 2018), in the second nursing period of these mice, mammary glands showed signs of premature involution at 15 days after delivery, when a significant increase of STAT-3 phosphorylation (p-STAT3, Y705) and cleaved caspase 3 (CC3) was determined (Figure 1A, B) and their litters resulted lighter than those from controls (Figure S1A). In addition, by hematoxylin-eosin (H&E) staining, we confirmed that mammary epithelium was underdeveloped in the first lactation of bi-transgenic compared with TTP^fl/fl^ mice, but impairment was even greater in the second reproductive cycle (Figure 1C). Furthermore, primary cultures of post-involuting mammary glands from MG TTP-KO females took longer to attach and grow than cells from control mice (Figure S1B), and when plated on ultra-low attachment plates, they formed less mammospheres than their control counterparts (Figure 1D). These data suggested to us that WAP-expresser PI-MECs may have been negatively affected by TTP loss.

To verify whether mammary progenitor compartments may be altered in other mouse models, we implanted mammary tissue fragments from total TTP-KO or C57BL/6 wild type (control) mice into inguinal cleared fat pads of 3-week-old syngeneic females. After 10-12 weeks, whole mount analysis revealed that mammary glands from implanted TTP-KO mammary epithelial cells developed significantly lower number and length of mammary tertiary ducts (Figure 1E). Similarly, mammary glands of 8 weeks old MMTV-Cre/TTP^fl/fl^ females displayed fewer branching ducts and a higher number of remaining TEBs than TTP^fl/fl^ control mice (Figure S1C).

**Figure 1.**
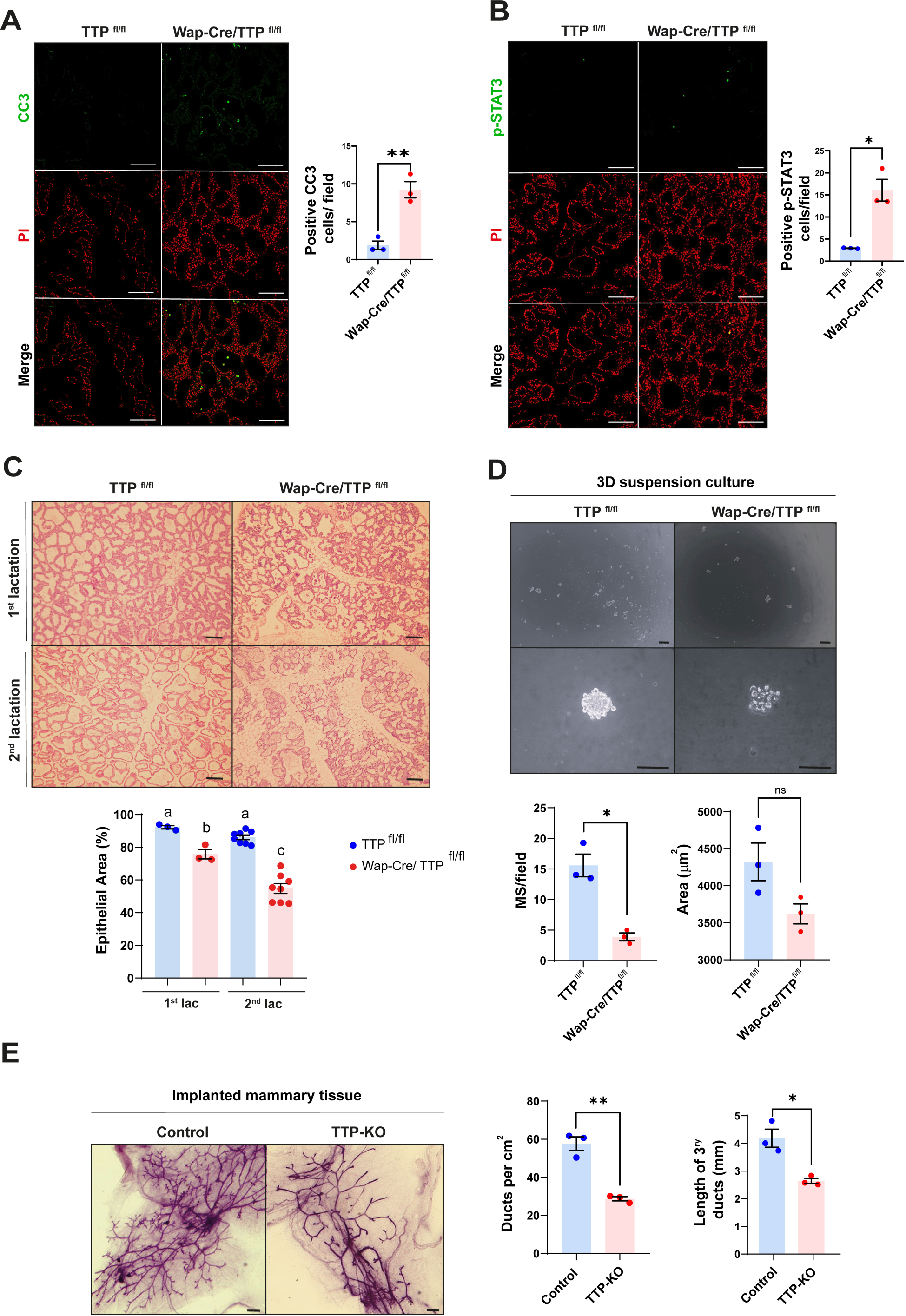
Phenotype of mammary glands of MG TTP-KO and total TTP-KO mice. **A & B.** Representative images of immunofluorescence (IF) assays for CC3 (A) and for p-Stat3 (B) of Wap-Cre/TTP^fl/fl^ (MG TTP-KO) females and TTP^fl/fl^ (control) mice at day 15 of their second lactation. Nuclei were stained with propidium iodide (PI). Corresponding quantification plots of positive nuclei per field ± SEM are shown on the right (T-Student test, n=3, *<0.05, **<0.01). Scale bar=100µm. OM=200X **C.** Representative images of H&E-stained mammary glands from TTP^fl/fl^ and Wap-Cre/TTP^fl/fl^ (MG TTP-KO) during first and second lactation. OM=100X. Scale bars=100µm. Below, quantification plots show % of area occupied by epithelium ± SEM, (One-way ANOVA and Tukey test, n= 3-8, *p<0.05). Significant differences exist between groups with no shared letters. **D.** Representative light field images of mammospheres (MS) at 10 days post-seeding derived from MECsof post-involuting TTP^fl/fl^ and Wap-Cre/TTP^fl/fl^ mammary glands (upper images: OM=40X;lower images: OM = 400X; scale bar = 100 µm). Below, quantification plots of average number of MS per field ± SEM and average area occupied by each MS ± SEM (T-Student test, *p<0.05, ns=non-significant). **E.** Representative images of whole mounted mammary glands from TTP-KO or control (C57BL/6 WT mice) implants into cleared fat pads of syngeneic female mice. Scale bars = 1mm. On the right quantification plots of average ductsper cm^2^ ± SEM and tertiary ductś length ± SEM (T-Student test, n=3, *p<0.05, ** p<0.01). OM: Original Magnification.

If TTP is relevant for mammary progenitor cell survival, one might expect higher *Zfp36* expression in at least some of these populations compared with other more differentiated ones. To test this hypothesis, we used our integrated single-cell RNAseq data set of post-natal mouse mammary glands (García Solá et al., 2021). This analysis showed that relatively high levels of TTP/*Zfp36* mRNA was detected in Basal cell clusters, *Procr*+ mammary stem cells and *Aldh1a3*+ luminal progenitor cells (Figure 2A). A similar distribution in the assembled UMAP was determined for all members of the Zfp36 gene network we have previously reported (Canzoneri et al., 2020) (Figure S2). Furthermore, focusing specifically on the luminal alveolar populations, we found relevant *TTP/Zfp36* expression levels mostly in progenitor cell clusters (Figure 2B).

**Figure 2:**
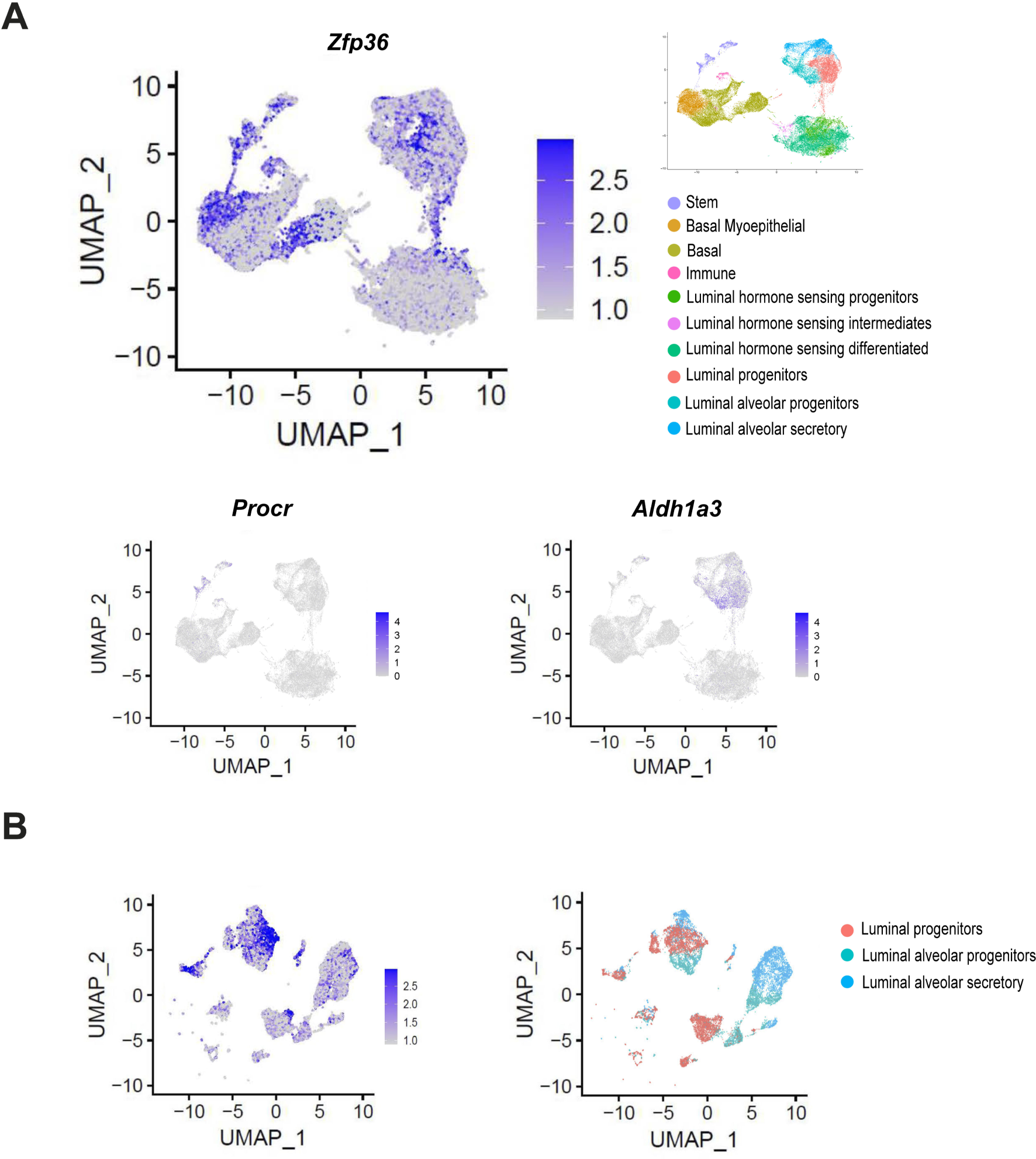
TTP/*Zfp36* mRNA expression analysis by integration of scRNAseq data sets in post-natal mouse mammary epithelial cells (MECs). **A.** Uniform Manifold Approximation and Projection (UMAP) plot of post-natal MEC populations displaying scRNAseq relative expression levels of *Zfp36, Procr and Aldh1a3* in a blue color intensity scale. **B.** UMAP plot of mammary luminal cell populations displaying relative expression levels of *Zfp36,* in a blue color intensity scale. Cell populations, as previously identified by García Solá et al, 2021, are depicted by colors in each panel.

In the human breast, most cancer subtypes have shown reduced *TTP/ZFP36* expression compared with normal tissue or “Normal-Like” mammary tumors (Canzoneri et al., 2020; Goddio et al., 2012). However, “Claudin-Low” breast cancers, which present a stem-like expression profile (Fougner et al., 2020), displayed significantly higher ZFP36 expression than the other subtypes, similar to the “Normal-like” levels (Figure 3A). Furthermore, using METABRIC publicly available data, we found positive correlations between *ZFP36* expression and genes associated with the stem-like phenotype, as *TWIST1, TWIST2, SNAI1, ZEB1, ZEB2, ALDH1A and YAP1* (Fougner et al., 2020) in all breast cancer subtypes (Figure 3B). Similar observations were made analyzing the TCGA-Pan Cancer data (Figure S3A, B) and it was determined that positive correlations of *TTP/ZFP36* expression with those stem-like markers were more significant in primary tumors (all subtypes considered) than in normal tissue (Figure 3C). Besides, analysis of this data set also revealed that in spite the clear TTP/*ZFP36* expression down-regulation in mammary tumors, there was not a better prognosis associated to high expression of this gene considering all breast cancer subtypes, and that free-disease survival time was even worse if only patients with basal-like subtype were considered (Figure 3D).

**Figure 3:**
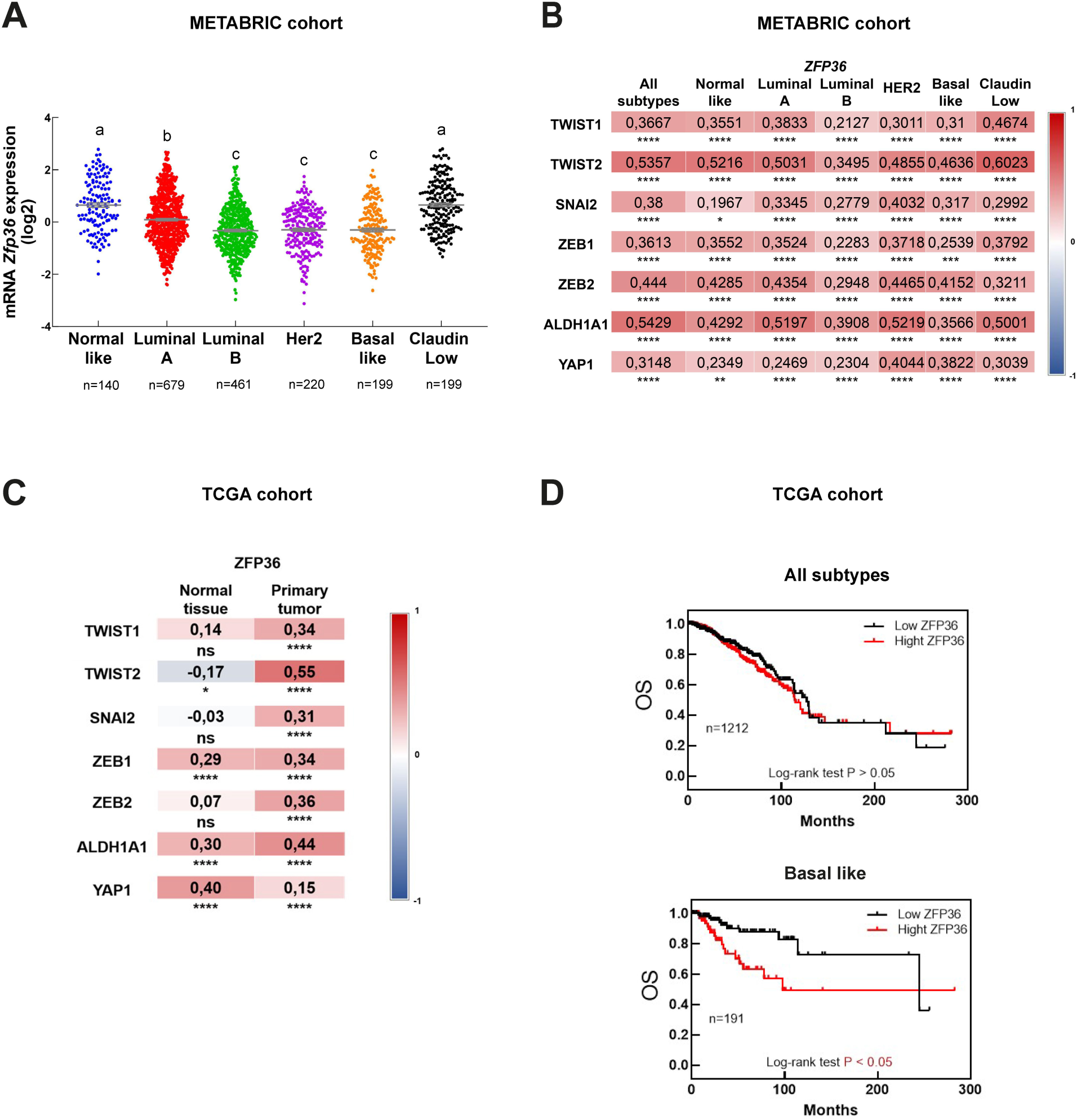
TTP/*ZFP36* mRNA expression in human breast cancer. **A.** Expression of *ZFP36* mRNA (log2 values) in breast cancer molecular subtypes according to METABRIC database. Kruskal Wallis and Dunńs tests were performed. Significant differences exist between groups with no shared letters (p<0.05). Number of tumors (n) is indicated below each plot. **B&C.** Expression correlation between *ZFP36* and genes related to stem/progenitor behavior comparing different breast cancer subtypes according to METABRIC database (B), and between normal tissue and primary tumors according to TCGA data (C). On the right of each graph, color scale (red: positive, blue: negative) indicates correlation level. Spearmańs ranked correlation test was applied (*p<0.05, **p<0.01, ***p<0.001, ****p<0.0001). **D.** Overall (OS) survival of breast cancer patients (all subtypes, on the left; basal-like subtype, on the right) from TCGA database categorized according to *ZFP36* expression levels. Number of patients (n) is indicated in each graph. To test significant differences, Log rank<0.05 was used.

### *TTP/ZFP36* knock-down decreases survival, self-renewal and repopulation capacity of HC11 mammary stem-like cells

To prove the impact of diminishing TTP levels in mammary progenitor cells, we proceeded to partly silence *Zfp36* by transfecting specific shRNAs complementary to different protein-coding regions in the HC11 mammary cell line, which display stem-like features (Ferrari et al., 2015; Sornapudi et al., 2018). Most clones in which TTP/*ZFP36* expression was significantly decreased did not survive more than a few passages. After evaluation of *TTP/Zfp36* expression levels on the clones that endured, we continue using the one transfected with shRNA 325.1(TTP-KD subline). These cells showed significantly less *Zfp36* mRNA and TTP protein expression than cells transfected with a *scrambled* shRNA (Sh-Ctrl cells) (Figure 4 A, B and Figure S4A).

**Figure. 4.**
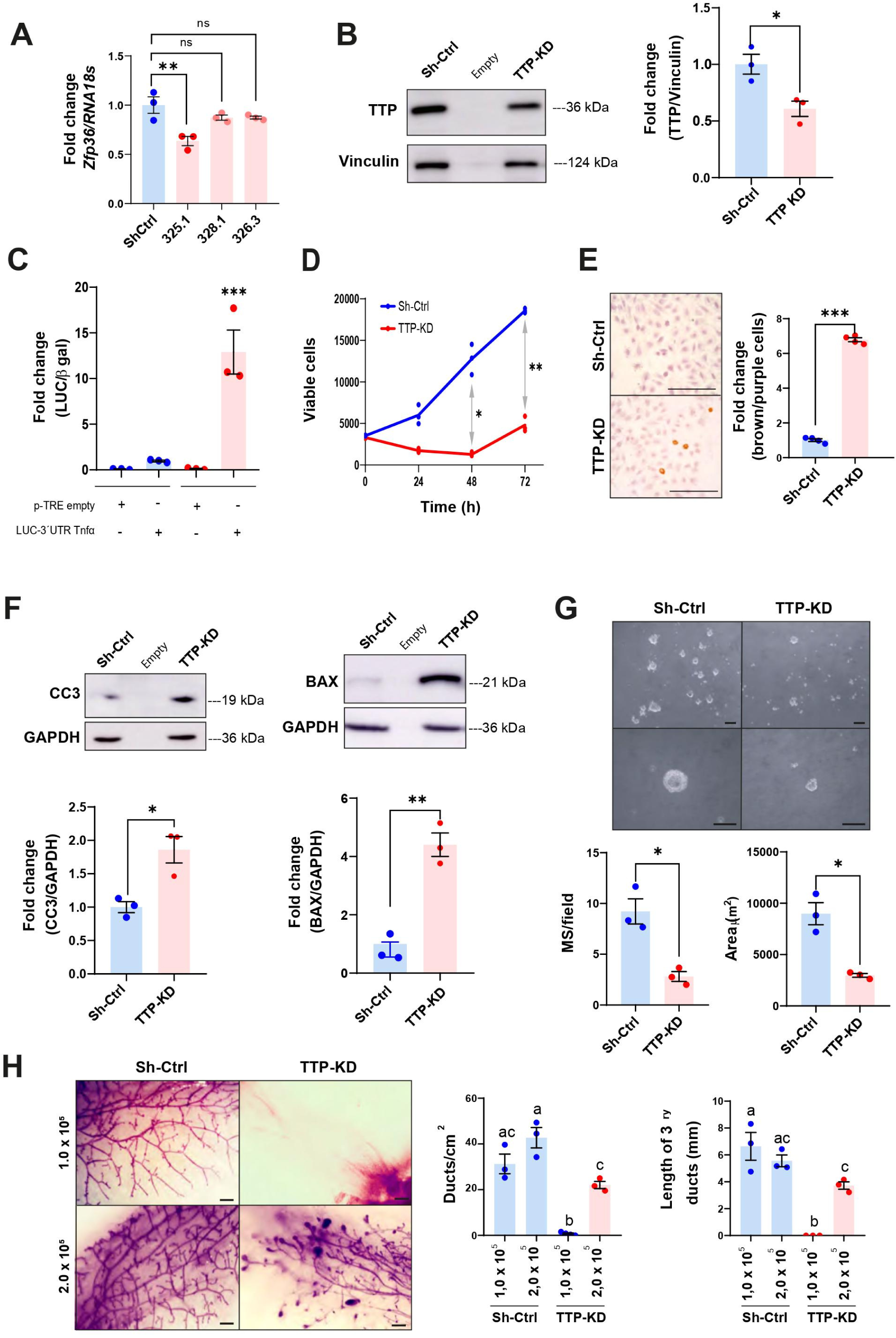
TTP/*Zfp36* knock-down decreases survival and self-renewal of HC11 mammary stem-like cells. **A.** *Zfp36* mRNA expression levels by RT-qPCR in HC11 clones stably transfected with shRNAs complementary to different *Zfp36* coding regions relative to HC11 cells transfected with a scrambled ShRNA (Sh-Ctrl cells). Values are shown as fold change of *Zfp36/RNA18s* ratio ± SEM (One way ANOVA, Dunnet test, n=3, ns=non-significant, **p<0.01). **B.** Representative image of TTP/*ZFP36* WB analysis in TTP-KD and Sh-Ctrl cells (left panel). On the right, quantification plot shows fold change of TTP/Vinculin protein expression ± SEM (T-Student test, n=3, *p<0.05). **C.** Luciferase activity in Sh-Ctrl and TTP-KD cells transfected with pTRE-LUC-3′UTR-*Tnfα* or pTRE empty vector plasmid and CMV-β-Galactosidase transfection control expression plasmid. Plotshows fold changes of Luciferase/β-Gal units ± SEM relative to Sh-Ctrl cells transfected with pTRE-LUC-3′UTR*-Tnfα* vector (ANOVA and Tukey test, n=3, ***p<0.001). **D.** Metabolic active Sh-Ctrl and TTP-KD cells analyzed by MTS every 24 h for 3 days. Plotshows average number of viable cells ± SEM (two-way ANOVA and Tukey test, n=3, *p<0.05, **p<0.01).Significant differences exist between groups with no shared letters . **E.** Representative images of Sh-Ctrl and TTP-KD cells analyzed by TUNEL assay. Cells with brown nuclei were considered apoptotic. On the right, plot shows fold change of apoptotic relatively to non-apoptotic cells ± SEM (T-Student test, n=4, ***p<0.001). **F.** Representative WB analysis of cleaved caspase 3 (CC3) and BAX in Sh-Ctrl and TTP-KD cells. Below, quantification plots showing protein fold changes of CC3/GAPDH ± SEM and BAX/GAPDH ± SEM in TTP-KD relatively to Sh-Ctrl cells (T-Student tests, n=3, *p <0.05, **p<0.01). **G)** Brightfield images of Sh-Ctrl and TTP-KD derived MS 10 days after seeding. OM = 100X (upper images), 400X (lower images). Scale bars=100µm. Below: plots showing average number of MS per field ± SEM and average MS diameter ± SEM in Sh-Ctrl and TTP-KD cells (T-Student test, n=3, *p<0.05). **H.** Representative images of whole mounted Sh-Ctrl or TTP-KD 1.0 x 10^5^ or 2.0 x 10^5^ cell implants in mammary cleared fat pads (scale bars=1mm). On the right, quantification plots of ductsper cm^2^ ± SEM and tertiary ductślength ± SEM (one-way ANOVA and Tukey test, p<0.05, n=3 implanted gland per condition). Significant differences exist between groups with no shared letters . OM: original magnification; MS: mammospheres.

To determine TTP activity in Sh-Ctrl and TTP-KD cells, they were transfected with the pTRE-LUC-3′UTR-*Tnfα* plasmid in which the TTP binding region of *Tnfα* 3′UTR has been located downstream Luciferase cDNA (Beisang and Bohjanen, 2012; Carballo et al., 1998). Figure 4C shows that luciferase activity increases about 12-fold in TTP-KD compared with Sh-Ctrl cells, although reduction in *TTP/Zfp36* expression is only around 40% (Figure 4 A, B).

TTP-KD cell line showed reduced viability in different culture conditions (Figure 4D and Figure S4B), increased apoptosis (Figure 4E), increased levels of pro-apoptotic proteins as BAX, cleaved caspase 3 (CC3) (Figure 4F) and impaired self-renewal capacity, since they produced smaller and fewer mammospheres than Sh-Ctrl cells in low-attachment plates (Figure 4G). Furthermore, when 1×10^5^ TTP-KD cells were implanted into BALB/c inguinal cleared fat pads, no mammary development was observed after 10 weeks, although Sh-Ctrl cells gave rise into fully developed glands. On the other hand, when the number of implanted cells was doubled, both HC11 sublines originated a normal mammary ductal network. However, the whole mount of glands originated from 2×10^5^ TTP-KD cells displayed ducts that do not reached the limits of the mammary fat pad and remaining TEBs, which have already disappeared in glands from Sh-Ctrl cells. This indicates that even increasing the number of implanted cells, TTP down-regulation results in a slower and/or incomplete ductal development possibly due to the impairment or depletion of the mammary progenitor population (Figure 4H).

### *TTP/ZFP36* down-regulation leads to inflammatory cytokine over-expression and activation of stress-associated signaling pathways

TTP-KD cells showed increase of *II-6* and *Tnfα* mRNA levels and phosphorylation of transcription factors commonly activated by these cytokines, as STAT3 and NF*k*B p65/RelA, respectively (Berishaj et al., 2007; Kalliolias and Ivashkiv, 2016) compared with Sh-Ctrl cells (Figure 5A,B). In fact, p65/RelA phosphorylation on S536 would be mostly dependent on TNFα overexpression in these cells, since adding the soluble receptor Etanercept was sufficient to block this effect (Figure 5C). Importantly, increase of p-p65 was also observed in the mammary glands of lactating MG TTP-KO compared with control mice (Figure 5D).

**Figure 5.**
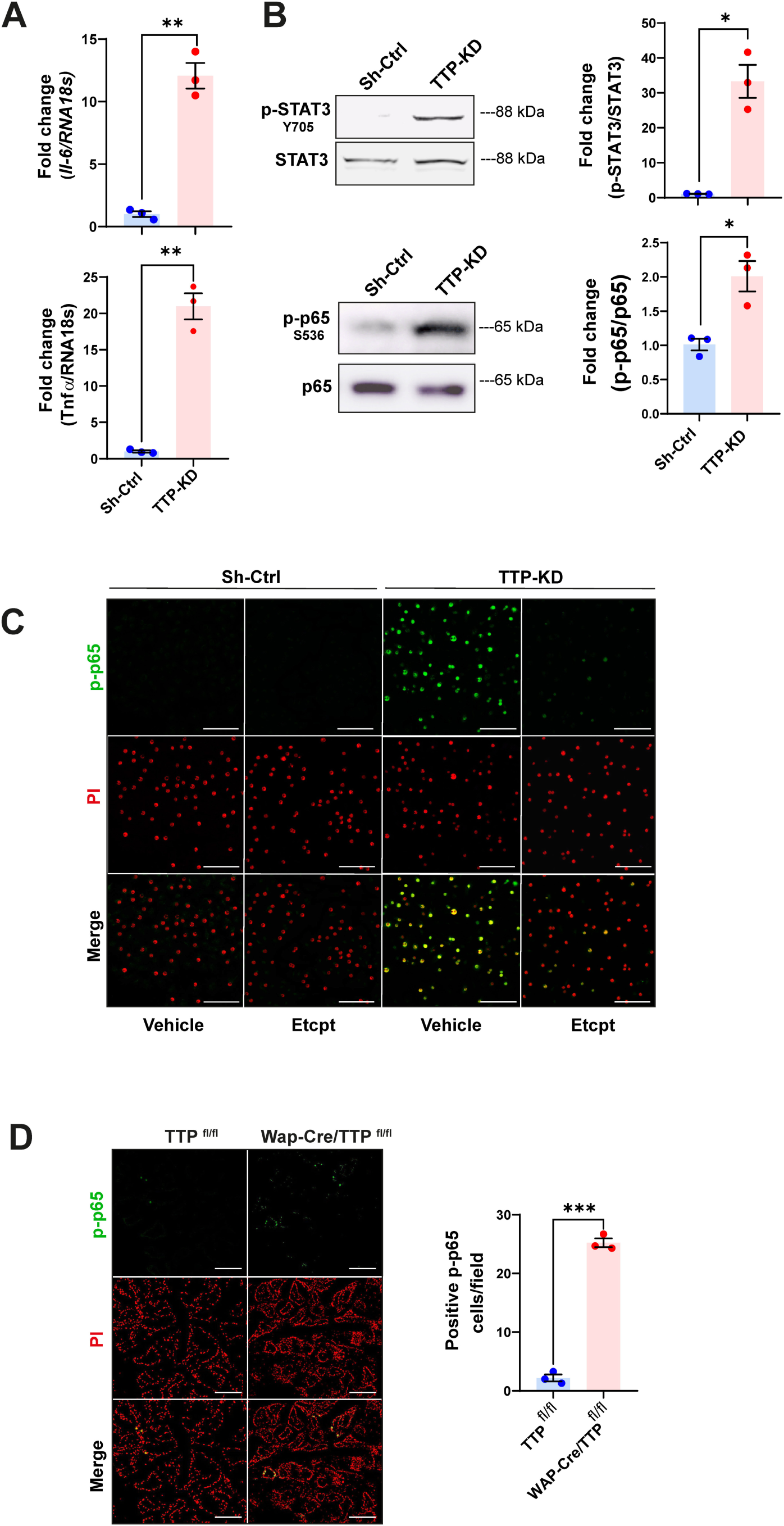
TTP/*Zfp36* down-regulation leads to activation of STAT3 and NFκB pathways in mammary epithelial cells. **A.** *Tnfα* and *Il-6 mRNA expression levels normalized to RNA18s* mRNA and showed as fold change ± SEM in TTP-KD relatively to Sh-ctrl cells (T-Student test, n=3, **p<0.01). **B.** Representative WB analyses of p-STAT3 (T705) and p-p65 (S 536) in Sh-Ctrl and TTP-KD cells (left panel). On the right, quantification plots showing fold change of p-STAT3/total STAT3 and p-p65/total p65 ± SEM in TTP-KD relatively to Sh-ctrl cells (T-student test, n=3, *p<0.05). **C.** Representative images of IF assays for p-p65 (S536) detection in Sh-Ctrl or TTP-KD cells treated with Etanercept or vehicle. Nuclei were labeled with Propidium Iodide (PI), after treatment with RNase. Original magnification=200X. Scale bars=100 µm. **D.** Representative images of IF analysis for p-p65 (S536) detection in Wap-Cre/TTP^fl/fl^ and TTP^fl/fl^ mammary glands at day 15 of their second lactation. Nuclei were labeled with Propidium Iodide (PI). On the right, quantification of p-p65 positive cells per field ± SEM (T-Student test, n=3, ***p<0.001). IF: immunofluorescence.

Figure 6A shows that when the MAPK phosphorylation pattern was analyzed, we found a drastic increase of constitutive p38 phosphorylation (p-p38, T183/Y182) in TTP-KD compared to control cells. On the other hand, ERK1/2 and JNK1/2 phosphorylation (Y204 and T183/Y185 respectively) was lower in TTP-KD compared with Sh-Ctrl cells. Interestingly, inhibition of p38, using SB203580, caused a rise of ERK1/2 and JNK1/2 phosphorylation, indicating that p38 overactivation may be responsible for ERK1/2 and JNK1/2 down-regulation in TTP-KD cells. On the other hand, ERK and JNK inhibition (by PD98059 and SP600125 treatment, respectively) decreased p-p38 levels in this cell line. Whether that was due to unspecific effects of these pharmacological inhibitors on p38 phosphorylation or to ERK1/2 and JNK 1/2 positive contribution to p38 phosphorylation remains to be elucidated.

**Figure 6.**
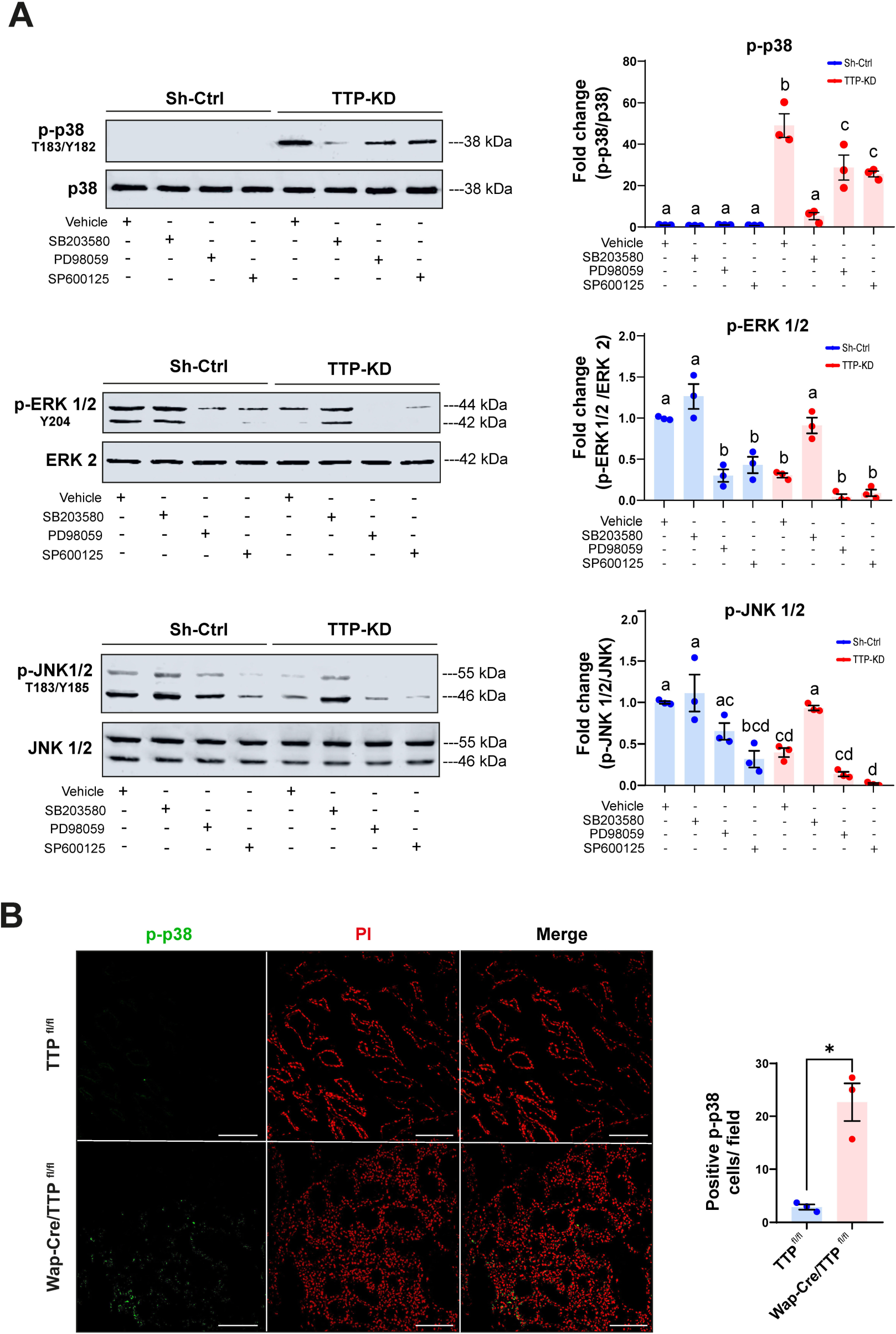
Regulation of MAPK activation in TTP-KD cells. **A.** Representative WB analyses of p-p38 (T180/Y182), p-ERK1/2 (Y204) and p-JNK1/2 (T183/Y185) in Sh-Ctrl or TTP/KD cells treated with either vehicle, 10μM p38 α and β inhibitor SB203580, 20μM MEK 1/2 inhibitor PD98059, , or 10 μM JNK 1/2 inhibitor SP600125. On the right, quantification plots show fold change of phosphorylated MAPKs/total MAPKs ± SEM in cells with different treatments relatively to Sh-Ctrl cells treated with vehicle (One way ANOVA and Tukey test, n=3, p<0.05). Significant differences exist between groups with no shared letters. **B.** Representative images of IF analysis for p-p38 (T180/Y182) detection in Wap-Cre/TTP^fl/fl^ and TTP^fl/fl^ mammary glands at day 15 of their second lactation. Nuclei were labeled with Propidium Iodide (PI). On the right, quantification of p-p38 positive cells per field ± SEM (T-Student test, n=3, *p<0.05). IF: immunofluorescence.

The over-activation of p38 as a consequence of TTP down-regulation was also evident in the mammary glands of lactating MG-TTP KO females, in which nuclear and cytoplasmic p-p38 labeling was observed. Nuclear localization was prioritized for quantification since it may be related with the pro-apoptotic activity of this enzyme (Cuadrado and Nebreda, 2010; Wood et al., 2009) (Figure 6B).

We determined that secreted TNFα is not only required for constitutive p65/RelA activation, but also for p38 phosphorylation in TTP-KD cells. Besides, interaction of the cytokine with its receptors as well as p38 activation is necessary for basal caspase 3 activation in this cell line (Figure 7A). Besides, Etanercept as well as SB203580 treatment prevented *Tnfα*-3’UTR stabilization induced by TTP down-regulation (Figure 7B). Finally, both reagents not only reversed, but greatly increased mammosphere formation capacity of TTP-KD cells (Figure 7C). This clearly shows TTP fundamental role protecting mammary progenitor cells from the deleterious consequences of TNFα-pp38 signaling pathway activation that would spontaneously occur if that AUBP is not fully active.

**Figure 7.**
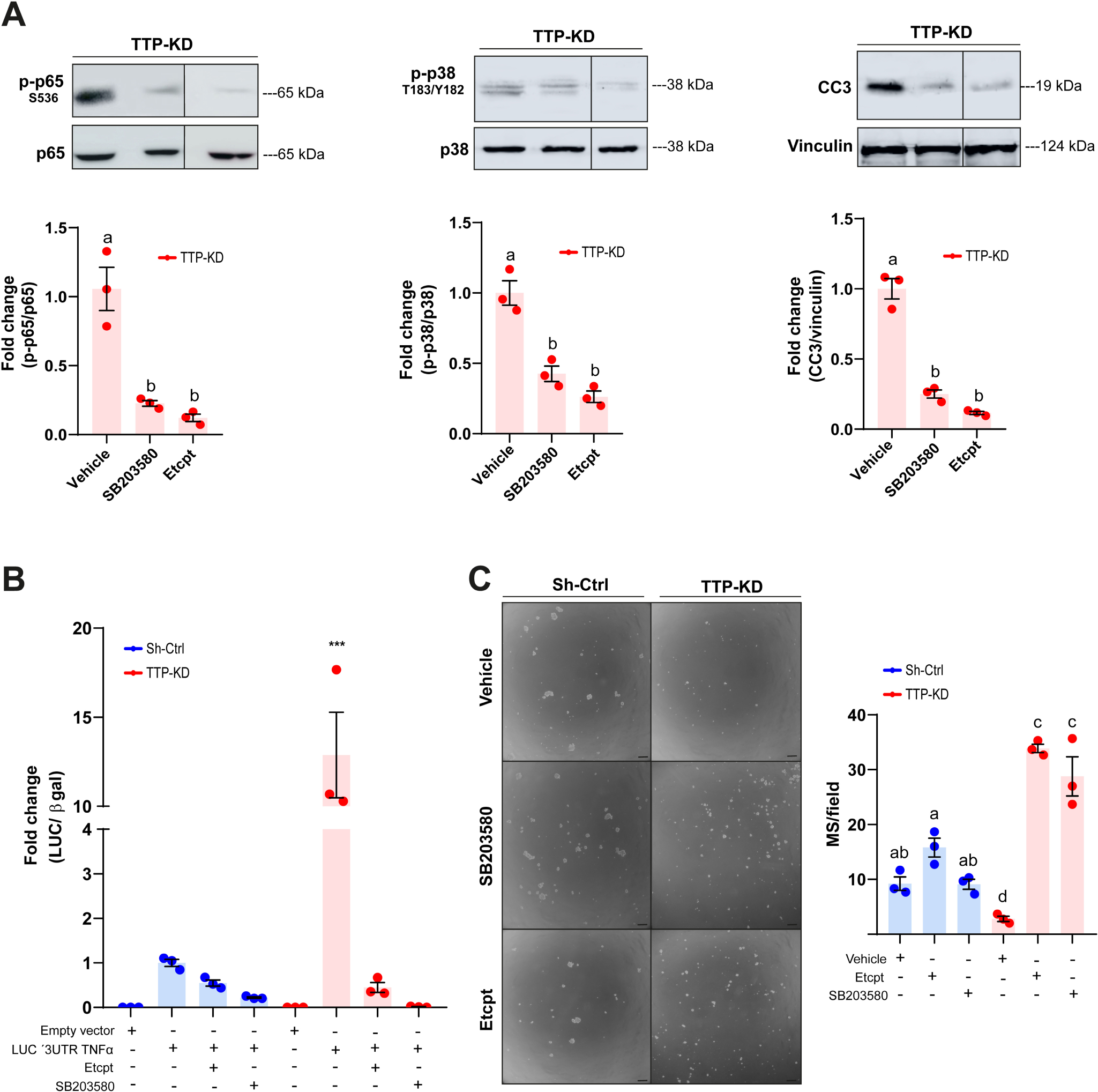
Inhibition of p-38 activation or TNFα blockade restores TTP activity and self-renewal in TTP-KD cells. **A.** Representative WB analyses of p-p65 (S536), p-p38 (T180/Y182) and CC3, normalized to total p38, total p65 and Vinculin, respectively in TTP-KD cells treated with vehicle, SB203580, or Etanercept. Below, quantification plots show fold change ± SEM relative to TTP-KD cells treated with vehicle. Significant differences exist between groups with no shared letters. **B.** Luciferase activity detected in Sh-Ctrl and TTP-KD cells transfected with pTRE-LUC-3′UTR-*Tnf*α or pTRE empty vector plasmid and CMV-β-Galactosidase transfection control expression plasmid and treated with 0.03 μg/μL Etanercept, 10 µM SB203580 or vehicle. Plotshows fold changes of Luciferase/β-Gal units ± SEM relative to Sh-Ctrl cells transfected with pTRE-LUC-3′UTR-*Tnfα* vector (ANOVA and Tukey test, n=3, ***p<0.001). **C.** Representative images of MS derived from Sh-Ctrl and TTP-KD cells treated with 0.03 μg/μL Etanercept, 10 µM SB203580, or vehicle for 10 days in 3D suspension culture. OM=100X. Scale bar=100 µm. On the right, plot shows average number of MS per field ± SEM (One way ANOVA and Tukey contrast, n=3, p<0.05). Significant differences exist between groups with no shared letters. MS: mammospheres. OM: original magnification.

## DISCUSSION

Here, we show that TTP*/Zfp36* expression would play a relevant role in the mouse mammary gland by contributing to the maintenance of different mammary progenitor cell populations. Bipotent MaSCs as well as long-lived unipotent cells in addition to diverse luminal progenitor subtypes have been identified in mouse and human mammary tissue (Visvader and Stingl, 2014). Particularly, the MG TTP-KO model allowed us to determine the importance of this protein in PI-MECs, which behave as multipotent mammary epithelial progenitors upon transplantation and as cancer-initiating cells in MMTV-Her2/neu multiparous transgenic mice (reviewed by Wagner and Smith 2005). They are able to form mammospheres in culture and express markers associated with mammary stem cells (Matulka et al., 2007). We have previously shown that TTP/*Zfp36* expression is highly increased during lactation, mainly by prolactin, through Stat5A activation, similarly to various milk protein genes. In addition, we have demonstrated that during lactation, TTP wards off early involution by preventing the untimely increase of local inflammatory factors (Goddio et al., 2018). Although it has not been determined whether PI-MECs originate from cells expressing WAP that are not terminally differentiated, or whether they arise from differentiated cells that bypass apoptosis during involution or both (Wagner and Smith, 2005), it can be speculated that TTP/*Zfp36* expression decrease observed in involuting mammary glands (Goddio et al., 2012) would correspond to down-regulation in the differentiated cells, making them susceptible for dying, while PI-MECs would maintain relatively high expression levels, preventing the impairment of this stem-like compartment.

Our analysis of scRNA-seq data sets confirm the previously reported observation about the conserved co-expression of TTP/*Zfp36* with other early-responsive genes (Canzoneri et al., 2020; Gutierrez et al., 2022) which here we show that occurs, at least partly, in *Procr+* MaSCs and *Aldh1a3*+ luminal progenitor cells. We have also found expression of this set of genes in some populations of the basal compartment, which might correspond to an “obligatory transitional“ transcriptional state described by Gutierrez et al., 2022). Once again, TTP/*Zfp36* expression might be relevant to maintain these mammary cell populations throughout different periods of the post-natal female mouse life.

Several reports have shown that TTP/*Zfp36* down-regulation is associated with breast cancer progression and/or treatment resistance (Al-Souhibani et al., 2010; Brennan et al., 2009; Canzoneri et al., 2020; Goddio et al., 2012; Griseri et al., 2011; Marderosian et al., 2006; Upadhyay et al., 2013). However, as pointed out previously (Goddio et al., 2012; Griseri et al., 2011), the relevance of TTP/*Zfp36* mRNA level for breast cancer prognosis is debatable. In fact, using TCGA data, we did not find a negative correlation between high expression of this gene and breast cancer patient survival. Besides, we show that the association between the mammary stem-like phenotype and TTP/*Zfp36* expression can be observed not only in mouse mammary cells, but also in human breast cancer tissue. Interestingly, positive correlation of this gene with different stem-like gene markers was stronger in primary tumors than in breast samples from healthy women, which might be due to a higher proportion of cells that express these markers in neoplastic tissue than in normal mammary epithelium.

The significance of TTP/*Zfp36* expression in the survival of stem-like cells was confirmed by the loss of cleared-fat pad repopulation and mammosphere formation capacity of TTP-KD HC11 cell line. These cells showed a dramatic decrease in their ability to reduce *Tnf*α mRNA post-transcriptionally probably due to the constitutive activation of p38, which phosphorylates the MAPK-activated protein kinase 2 (MK2) that, in turn, would block the activity of the remnant expressed TTP/*Zfp36* (Tiedje et al., 2016).

Although TTP modulates multiple cytokines, it clearly exerts a dramatic effect on TNFα production by controlling its expression at transcriptional, post-transcriptional and translational level (reviewed by Guo et al., 2017). Furthermore, it has been shown that cachexia, arthritis, and autoimmunity developed by TTP-KO mice could be prevented if they were early treated with TNFα antibodies, indicating that the phenotype was mostly due to the excess of this cytokine (Taylor et al., 1996). We found that a pathway triggered by extracellular TNFα would be mostly responsible for apoptosis induction in the TTP-KD cells. Nevertheless, higher levels of IL-6 and hyper-activation of STAT3 might be also relevant for inducing cell death, as previously seen in involuting mammary glands (Hughes and Watson, 2018; Zhao et al., 2002). Similarly, in lactating MG-TTP KO female mice, which also displayed IL-6 over-expression and STAT3 phosphorylation, excessive production and secretion of TNFα played a key role in cell death induction (Goddio et al., 2018).

Our results show that in TTP-KD cells TNFαinduces p38 phosphorylation, as reported in other models (Kalliolias and Ivashkiv, 2016; Sabio and Davis, 2014), and that activation of this MAPK leads to mammary cell death, as it has also previously observed (Wen et al., 2011). Besides, constitutive over activation of p38 may be, at least partly, also responsible for down-regulation of ERK1/2 and JNK1/2 phosphorylation. This effect would be similar to what reported in macrophages by Hall and Davis, 2002.

In conclusion, our results indicate that expression of TTP/*Zfp36* in the mammary gland would be not only important for controlling expression of proteins that may contribute to tumor progression, for maintaining lactation by preventing spontaneous initiation of involution associated inflammatory pathways, but also for maintaining the survival of progenitor slow cycling cell populations that are fundamental for fueling development during puberty and also in each reproductive cycle of the female mouse.

## EXPERIMENTAL PROCEDURES

### Mouse models

Mice were maintained in a specific pathogen-free facility at the FCEN-UBA-CONICET, at constant temperature of 22°C ± 2°C and 40-70% humidity with a 12-hours light cycle. Animals were allowed food and *water ad libitum*. Mouse experiments were approved by local IACUC authorities and complied with regulatory standards of animal ethics. As previously reported (Goddio et al., 2018), to obtain the bi-transgenic Wap-Cre/TTP^fl/fl^ (MG TTP-KO) strain, we crossed TTP^fl/fl^ with Wap-Cre mice, both in C57BL/6 genetic background. TTP^fl/fl^ originated and gently provided by Dr. Perry Blackshear lab (NIEHS, NIH, USA), presents the exon 2 of *Zfp36,* which codes for the zinc finger domain, surrounded by LoxP sequences (Qiu et al., 2012). The Wap-Cre mice express Cre recombinase specifically in mammary gland epithelial cells driven by the mouse milk protein *Wap* (whey acidic protein) promoter. This strain was obtained from the NCI-NIH Mouse Repository. Genomic DNA was obtained from tail samples by a “HotSHOT” technique as previously described (Truett et al., 2000). Wap-Cre/TTP^flfl^ mice were identified by PCR using 150 ng of DNA per reaction. *Zfp36* floxed and *Zfp36* WT alleles were amplified with P1_Fw and P2_Rv primers. Wap-Cre allele was amplified with W003_Fw and C031_Rv primers (see **Suppl. Table 1**). The removal of *Zfp36* exon 2 in MG-TTP KO mammary glands was tested by multiplex PCR with two forward primers (P1_Fw, P3_Fw) and reverse primer P4_Rv. Product size of each PCR are shown in Suppl. Table 2 and corresponding agarose gel images are shown in **Figure S5**.

MMTV-Cre mice (from the NCI-NIH mouse repository) were crossed with TTP^fl/fl^ mice. Bitransgenic female mice were detected by PCR with the same method described for Wap-Cre/TTP^flfl^ mice. MMTV-Cre allele was amplified using MMTV-Cre_Fw and CRE_C031_RV primers (Suppl. Table 1). Product size is shown in Suppl. Table 2. KO of TTP was checked by the same multiplex PCR as explained above (Figure S5). Females of 8-week-old were euthanized and whole mount analysis of their mammary glands were performed.

Inguinal mammary fat pads of 21-day-old female C57BL/6 (for TTP-KO tissue implants) or BALB/c (for HC11 cell injection) mice were cleared of endogenous epithelium to perform implantation studies (Blair and DeOme, 1961) of total TTP-KO female mouse (Taylor et al., 1996) mammary fragments or HC11 TTP-KD or Sh-Ctrl, cell sublines. In the last case, suspensions of 1.0 x 10^5^ and 2.0 x 10^5^ cells of each subline were prepared in a way that the volume injected in each fat pad was not superior to 100 µL. After 10 weeks, recipient mice were euthanized and whole mounts of mammary glands were performed. Since in adults the tertiary ducts, which come from the second branching event, are the ones that actively grow in the fat pad (Huebner and Ewald, 2014; Macias and Hinck, 2012), they were measured in quantity and length.

### Monolayer cell culture, transfections and treatments

HC11 mouse mammary gland cell line and its derived sublines were cultured at 37°C and 5% CO_2_ in RPMI with HEPES (Gibco, #23400021) supplemented with 5 μg/ml insulin, 1% antibiotic antimycotic (Gibco, #15240062) and 10 % Fetal Bovine Serum (Internegocios, #FBI). The TTP-KD HC11 sub-line was initiated by transfecting *Zfp36* sh-RNAs (Mission-Merk TRCN0000238325, TRCN0000238326, TRCN0000238328) cloned into pKLO.1-Puro vectors. These vectors were transfected with PEI (Polysciences), using 5 μg of total DNA in 6-well plates, (PEI: DNA 3:1). Clones were selected with 3 μg/mL (and maintained with 1μg/mL) of Puromycin (Invivogen, #ant-pr-5b).

Plasmids pTRE2hygLUC-3′UTR-*Tnfα* (gently provided by Dr. Jonathan Lamarre, Ontario Veterinary College, University of Guelph, Canada) and pCMVLacZ were transfected with Lipofectamine 3000 (Thermo, #L3000001) using1 µL/µg DNA, P3000 Reagent at 2 µL/µg DNA, and opti-MEM (Gibco, #A4124801). To harvest cells and perform luciferase readings, Reporter Lysis Buffer (Promega, #E1531) was employed.

For inhibiting MAPKs p38, ERK1/2 and JNK1/2, 10 µM SB203580, 20 µM PD98059 and 10 µM SP600125 (Calbiochem #559389, #513000 and #420119) or vehicle (DMSO) were used in medium with no FBS for 2h.

### RT-qPCR assays

Monolayer cultures were harvested with RNA-PrepZOL (Inbiohighway, # R0010) according to manufacturer instructions. RNA was reversed transcribed (RT) using MMLV InbioHighway (#E1601) reagent. All qPCRs were performed using SYBER Green (Roche) in the StepOnePlus equipment (Thermo). Gene expression levels were normalized to *RNA 18s* using standard curves (sequences in **Suppl. Table 3**).

### Cell viability and apoptosis assays

We used the *CellTiter 96® AQ_ueous_ One Solution Cell Proliferation Assay* (Promega, #G3582) for determining the number of viable cells, according to manufacturer instructions. Calibration curves were made by adding MTS 24 h post-plating and measuring the absorbance at 490 nm between 2.5 and 3.5 h later. With the obtained regression equation, it was possible to calculate how many cells were viable every 24 h using 3000 trypan-blue negative cells at the starting point.

Apoptotic cells were detected using the *DeadEnd™ Colorimetric TUNEL System* (Promega, #G7360) to perform the terminal deoxynucleotidyl transferase-mediated dUTP nick-end labeling (TUNEL) assay following manufacturer instructions. Diaminobenzidine (DAB) was employed as chromogen and hematoxylin ascounterstaining. At least 4 fields per condition were photographed and quantified. For positive control, cells were treated with DNase I (4u/mL) for 10 minutes.

### Primary cultures

Female mice at 21 days post-weaning were euthanized by cervical dislocation under sterile conditions. Right and left mammary glands # 2, 3 and 4 (without inguinal ganglion) were obtained with sterilized surgery tools and mechanically dissected in RPMI supplemented with HEPES, penicillin/streptomycin 1% v/v, 0.15% collagenase type 4, 0.2% trypsin, DNase 10 µg/mL and 2% FBS (Fetal Bovine Serum). Then, they were incubated in agitation for 1 h at 37°C and centrifuged several times to remove adipocytes and keep the pellet of mammary epithelial cells, which were resuspended in RPMI supplemented with HEPES, penicillin/streptomycin 1% v/v and 10% FBS, plated and monitored for up to 4 days. When epithelial cells were attached, they were tripsinized and used to carry out mammosphere assays.

### Mammosphere assays

Ultra-Low Attachment 6-well plates (Corning, # CLS3471-24EA) were used. Cells from primary cultures were plated at low density (10000 cells/mL) in RPMI plus HEPES media supplemented with 1% antibiotic/antimycotic (Gibco #15240062), 5 μg/mL insulin, 20 ng/mL human recombinant EGF (Thermo #PHG0313) and 1X Gem21 NeuroPlex™ Serum-Free (B27) (GeminiBio, #400-160). No serum was added in these assays. In those involving TTP-KD or Sh-Ctrl cells, 1 μg/mL Puromycin was added. Mammosphere formation was assessed 10 days post-plating by taking photos of 5 to10 fields per condition and analyzed by ImageJ. MAPK p38 was inhibited and TNFα was blocked by adding either 10 µM SB203580 or 0.03 µg/µL Etanercept in 2% FBS-RPMI for 8-10 days.

### Western blot (WB) analysis

Cells were harvested in RIPA buffer with 1:100 protease inhibitor cocktail set I (Calbiochem, #539131) and phosphatase inhibitors (1:100 Na_2_Ov_4_; 1:20 NaF). Most membranes were blocked for 1 h in 5% milk, but those for detection of phosphorylated proteins were blocked overnight in 5% gel fish (Sigma, #G7765). Blots were probed with antibodies listed in Suppl. Table 4. For MAPK analysis, fluorescent secondary antibodies were used and detected with the *Odyssey* imaging system. For studying other proteins, horseradish peroxidase (HRP) conjugated secondary antibodies were utilized and detection was performed with *Amersham ImageQuant 800*. Original WB images are shown in Figure S6.

### Immunohistofluorescence

Mammary glands were fixed in 4% paraformaldehyde (PFA) included in *OCT Compound*, frozen, and sectioned in 20 µm slides by cryostat. For DNA staining, slides were incubated with RNase A (10 µg/mL) for 3 h at 37°C and 1µg/mL propidium iodide (PI) was added for 5 min. Primary and secondary antibodies are listed in Suppl. Table 5.

### Mammary whole mounts (WM)

Whole mammary glands were extended on slides and fixed for 2 h in Carnoýs solution (6:3:1 Ethanol:Clorophorm:Glatial acetyl acid). Tissue was hydrated to be stained O.N. with carmin-alum, dehydrated and mounted.

### In silico analysis

Analysis of gene expression throughout different molecular subtypes of breast cancer were performed using The Cancer Genome Atlas (TCGA) -Pan-Cancer database and the METABRIC dataset. To evaluate expression levels differences, Kruskal Wallis and Dunn tests were carried out. To determine correlations, Spearmańs ranked correlation test was used. For scRNA-seq analysis, matrices from Bach et al., 2017, Giraddi et al., 2018 and Pal et al., 2017 were downloaded and loaded into R, version 3.6.3. The data corresponding to adult mice mammary samples were integrated into a single matrix and analyzed using the Seurat R package, version 3.4.4. A UMAP was generated using the top 3,000 variable genes selected via the default Variance Stabilizing Transformation (VST) method. Cluster/cell type labels were preserved from the original manuscripts for visualization and downstream analysis. Specific details can be found in García Solá et al., 2021.

### Statistical tests

Statistical significance of differences was evaluated using the software GraphPad Prism8. Data was presented as mean ± SEM unless otherwise noted. In every case, at least 3 independent experiments were evaluated (n = 3 or more).

### Data and code availability

TCGA Pan-Cancer and METABRIC consultation dates were 06/27/2022 and 06/28/2022. Sc-RNAseq data used in this paper are deposited in the Gene Expression Omnibus (GEO) database under accession codes GSE106273 (Bach et al., 2017), GSE103275 (Pal et al., 2017) and GSE111113 (Giraddi et al., 2018).

## ACKNOWLEDGEMENTS

We would like to thank Dr. Perry Blackshear for gently providing us the TTP mouse models and the mouse TTP antibody, and Dr. Marcelo Rubinstein for his help in the establishment of the Wap-Cre and MMTV-Cre mouse colonies in our animal facility.

## AUTHOR CONTRIBUTIONS

Conception, design, supervision: Edith C. Kordon, Omar Coso, Micaela Stedile. Writing: Edith C. Kordon and Micaela Stedile. Wet lab techniques: Micaela Stedile and Inés Beckerman. Transgenic mice techniques: Micaela Stedile, Inés Beckerman, Angela Lara Montero, Maria Victoria Goddio, Marina Ayre, Emilia Bogni, and Zaira Naguila. In silico analysis: Martín García Solá and Angela Lara Montero.

## FIGURE LEGENDS

