## Supplementary data for "Tristetraprolin promotes survival of mammary progenitor cells by restraining TNFα levels"

### SUPPLEMENTARY FIGURE LEGENDS

**Figure S1**. **A.** Weight of pups at 15 days post-natal from TTP$^{fl/fl}$ or Wap-Cre/TTP$^{fl/fl}$ mothers going through second lactation (weight ± SEM, n = 10-13, ***p<0.05). **B.** Bright field images of primary cultures of mammary glands´ tissue from TTP$^{fl/fl}$ or Wap-Cre/TTP$^{fl/fl}$ mice at 20 post- weaning. Above: 3 days post-plating; below: 5 days post plating. OM=400X. Scale bars=100 µm. **C.** Representative images of whole mounted TTP$^{fl/fl}$ and MMTV-Cre/TTP$^{fl/fl}$ mammary glands of 8-10 nulliparous female mice. Scale bars=1 mm.

**Figure S2.** UMAP of post-natal MEC populations displaying scRNAseq relative expression levels of *Zfp36* **(left panels)** in a red color scale and *Egr1, Fos, Fosb, Jun, Junb* AP-1 related genes **(central panels)** in a green scale. **Right panels:** Co-expression blend analysis of *Zfp36* and each of the AP-1 related genes. Here yellow dots depict cells where both genes considered are co-expressed, as shown in the color reference.

**Figure S3**. **A.** Expression of *ZFP36* mRNA (log2 values) in breast cancer molecular subtypes according to TCGA database. Kruskal Wallis and Dunn´s tests were performed. Significant differences exist between groups with no shared letters (p<0.05). **B.** Expression correlation between *ZFP36* and genes related to stem/progenitor behavior comparing different breast cancer subtypes according to TCGA database. On the right of each graph, color scale (red: positive, blue: negative) indicates correlation level. Spearman´s ranked correlation test was applied (*p<0.05, **p<0.01, ***p<0.001, ****p<0.0001).

**Figure S4. A.** Representative images of IF analysis for TTP detection in Sh-Ctrl and TTP-KD cells. Nuclei were stained with propidium iodide (PI). OM=200X, scale bar=100 µm. **B.** Metabolic active Sh-Ctrl and TTP-KD cells analyzed by MTS every 24 h for 3 days using 10% charcoal stripped FBS media or 10% FBS media supplemented with 2,5 ng/mL Epithermal Growth Factor (EGF). Plot shows average number of viable cells ± SEM (two-way ANOVA and Tukey test, n=3, *p<0.05, **p<0.01). Significant differences exist between groups with no shared letters.

**Figure S5. A.** Representative agarose gels images of Wap-Cre (left), MMTV-Cre (center) and *Zfp36* (right) genotyping PCR products. Product size is showed on the right of each gel. Genotype of mice tested is indicated below each well. In all cases DNA from tails was used. **B.** Representative agarose gels of *Zfp36*-KO PCR products performed in MECs of control (*Zfp36*$^{fl/fl}$ mice) or TTP-KO mice (Wap-Cre/*Zfp36*$^{fl/fl}$ mice on the right gel and MMTV-Cre/*Zfp36*$^{fl/fl}$ mice on the left gel). Control mice exhibit an 870 pb band (*Zfp36* flox alleles) and TTP-KO mice exhibit a 769bp band (*Zfp36* recombined or Δ alleles*).* Primers used for all these PCRs are specified in Suppl. Table 1 and product sizes in Suppl. Table 2.

**Figure S6. A.** Original Vinculin and TTP WB images corresponding to main Figure 4B. **B.C.** Original GAPDH, CC3 and BAX WB images corresponding to main Figure 4F. **D.E.** Original pSTAT3/STAT3, p65 and p-p65s WB images correspondending to main Figure 5B. **F. G. H.** Original p-p38/p38, p-ERK1/2/ERK2 and pJNK1/2/JNK1/2 WB images correspondending to main Figure 6A. **I. J. K.** Original p65, p-p65, p-p38/p38, Vinculin and CC3 WB images corresponding to main Figure 7A. In all cases, the bands shown in main figures are surrounded by dotted squares. Blots of panels A, B, C, E, I and K were revealed with ECL and images taken with the equipment *Amersham ImageQuant 800.* Blots of panels D, F, G, H and J were incubated with fluorescent secondary antibodies, using the red channel for total proteins and the green channel for phosphorylated forms. Those images were taken with the equipment *Odissey*. Only to observe p-p65, membranes were stripped and re-revealed for p65.

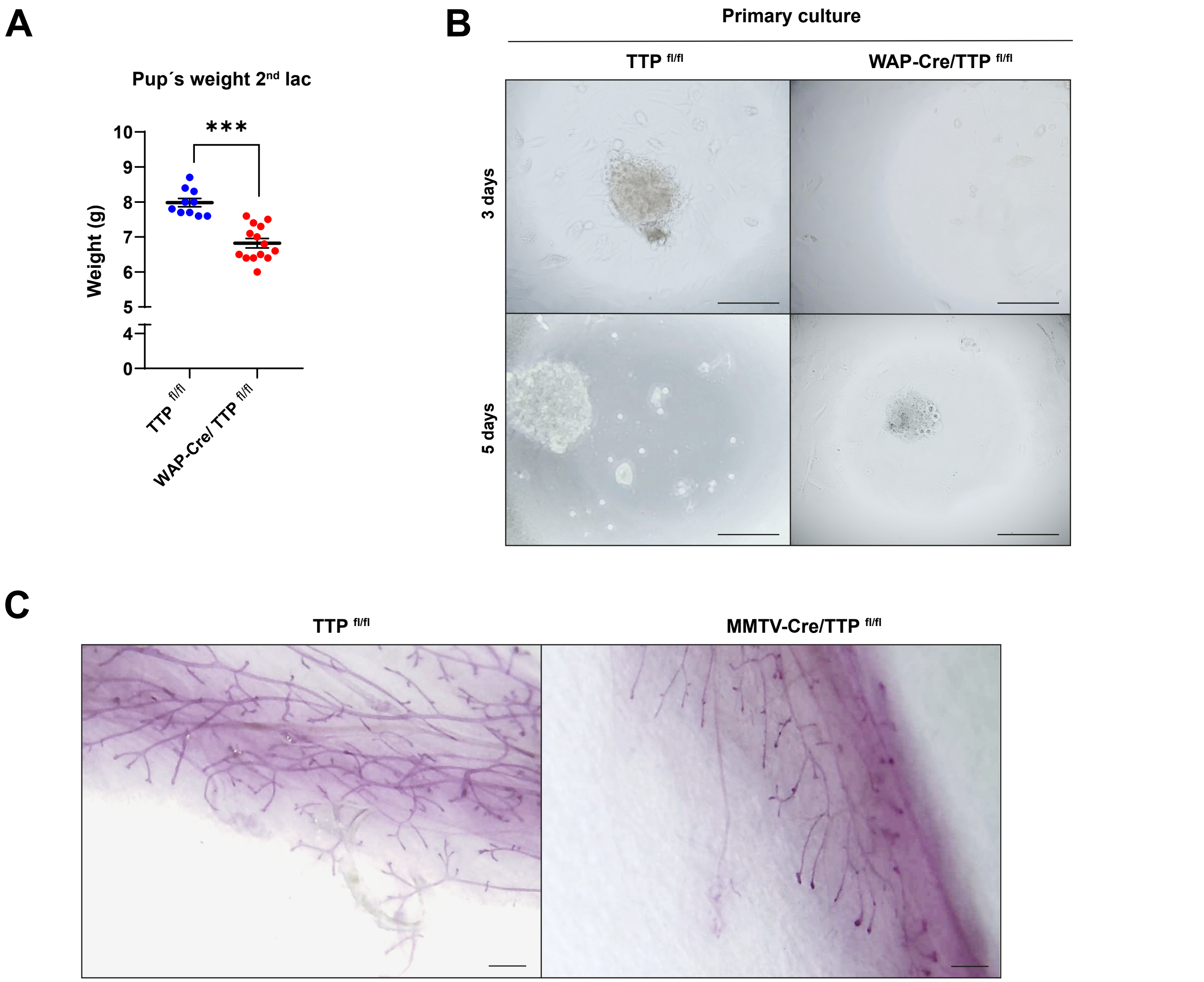


Supplementary Figure 1

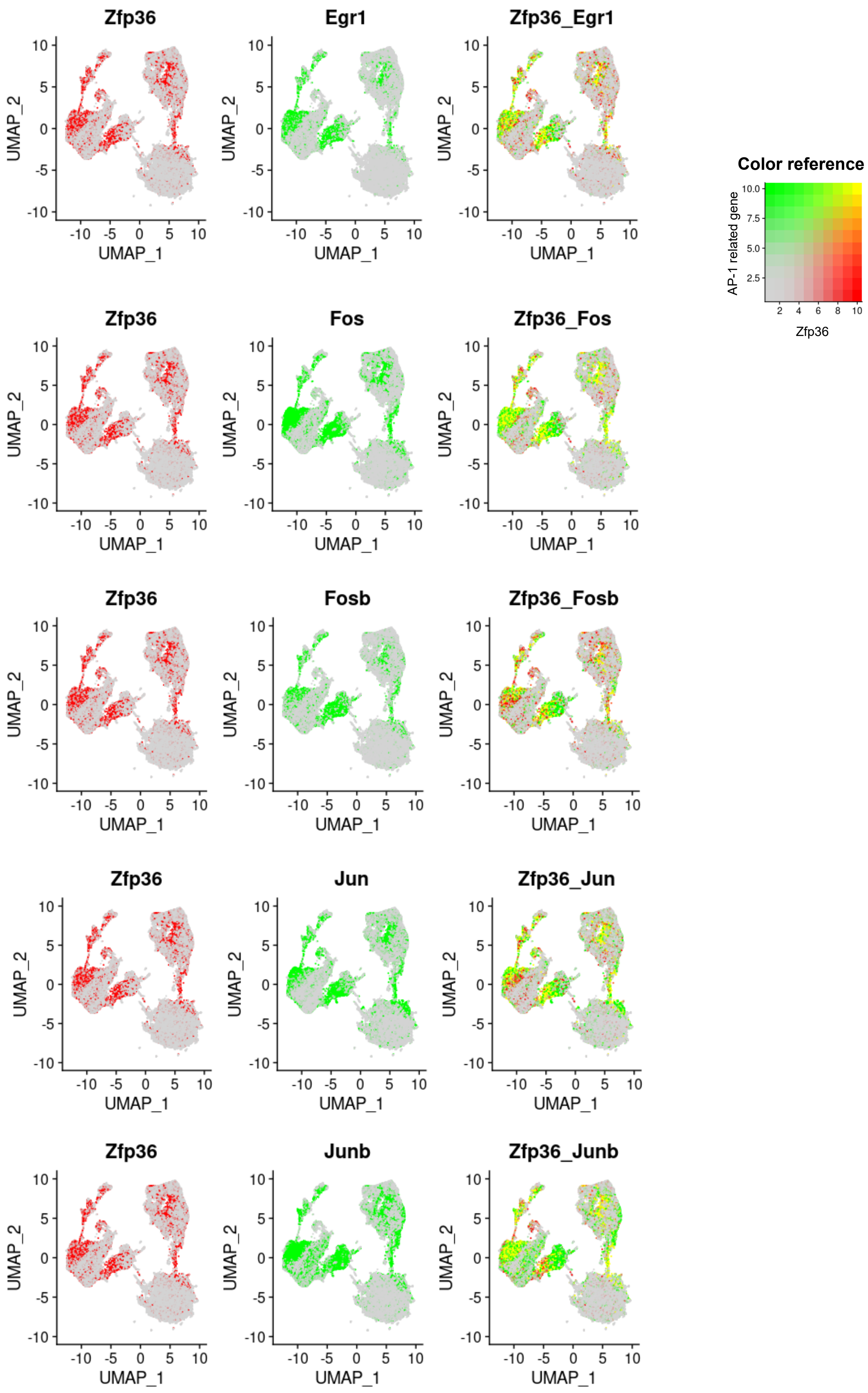


Supplementary Figure 2

**A**

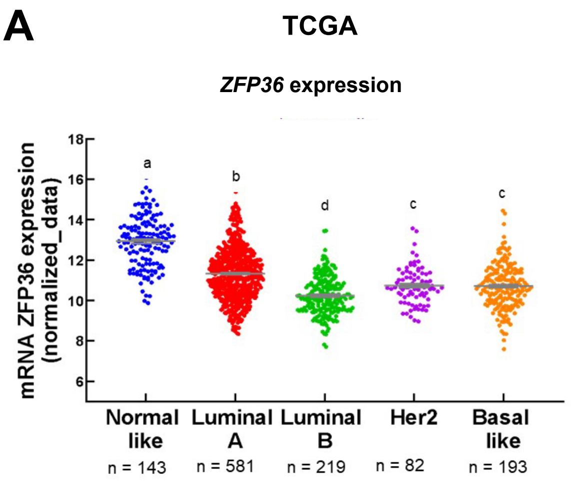


**B**

**TCGA**

| | ZFP36 | | | | | |
|---|---|---|---|---|---|---|
| | All subtypes | Normal like | Luminal A | Luminal B | HER2 | Basal like |
| TWIST1 | 0,43 **** | 0,43 **** | 0,43 **** | 0,09 ns | 0,36 *** | 0,23 ** |
| TWIST2 | 0,61 **** | 0,40 **** | 0,53 **** | 0,39 **** | 0,43 **** | 0,44 **** |
| SNAI1 | 0,38 **** | 0,36 **** | 0,39 **** | 0,33 **** | 0,23 * | 0,31 **** |
| ZEB1 | 0,41 **** | 0,30 *** | 0,33 **** | 0,20 ** | 0,08 ns | 0,24 *** |
| ZEB2 | 0,44 **** | 0,30 *** | 0,38 **** | 0,23 *** | 0,18 ns | 0,28 **** |
| ALDH1A1 | 0,53 **** | 0,40 **** | 0,51 **** | 0,19 ** | 0,27 * | 0,23 ** |
| YAP1 | 0,30 **** | 0,12 ns | 0,18 **** | 0,14 * | 0,00 ns | -0,16 * |

Color scale: 1, 0, -1

**Supplementary Figure 3**

A

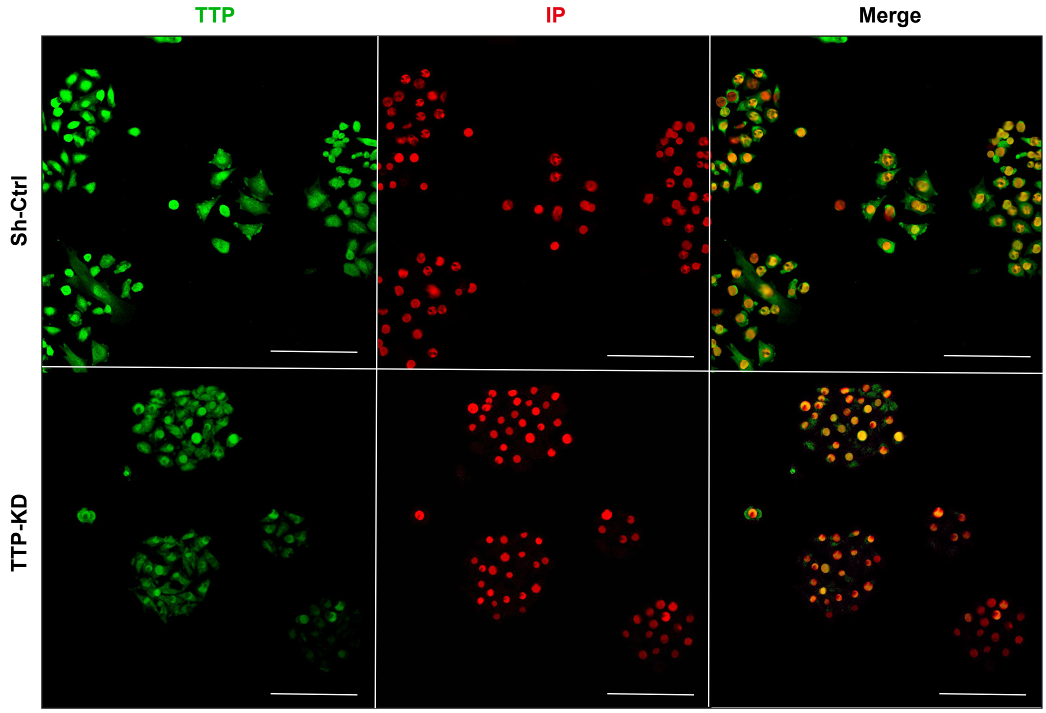


B

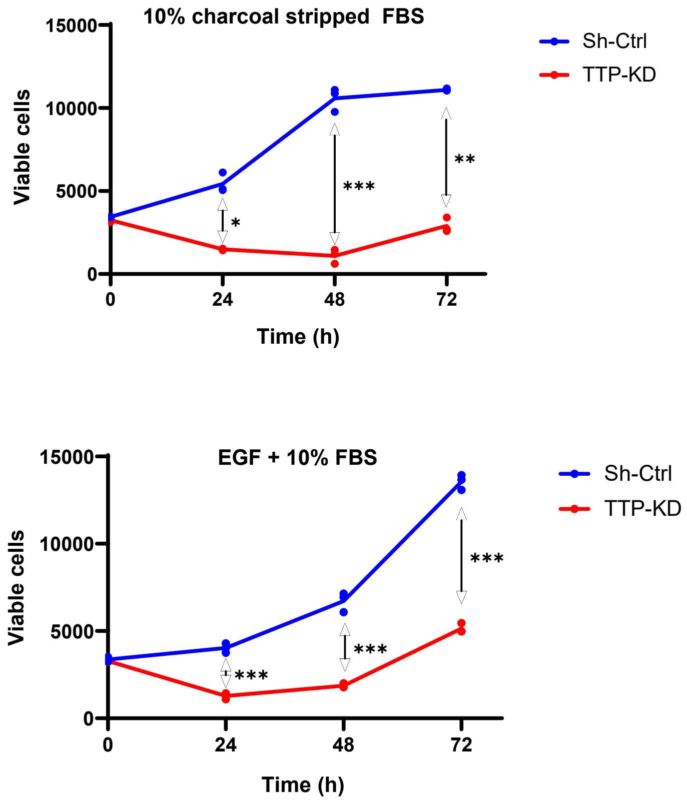


**Supplementary Figure 4**

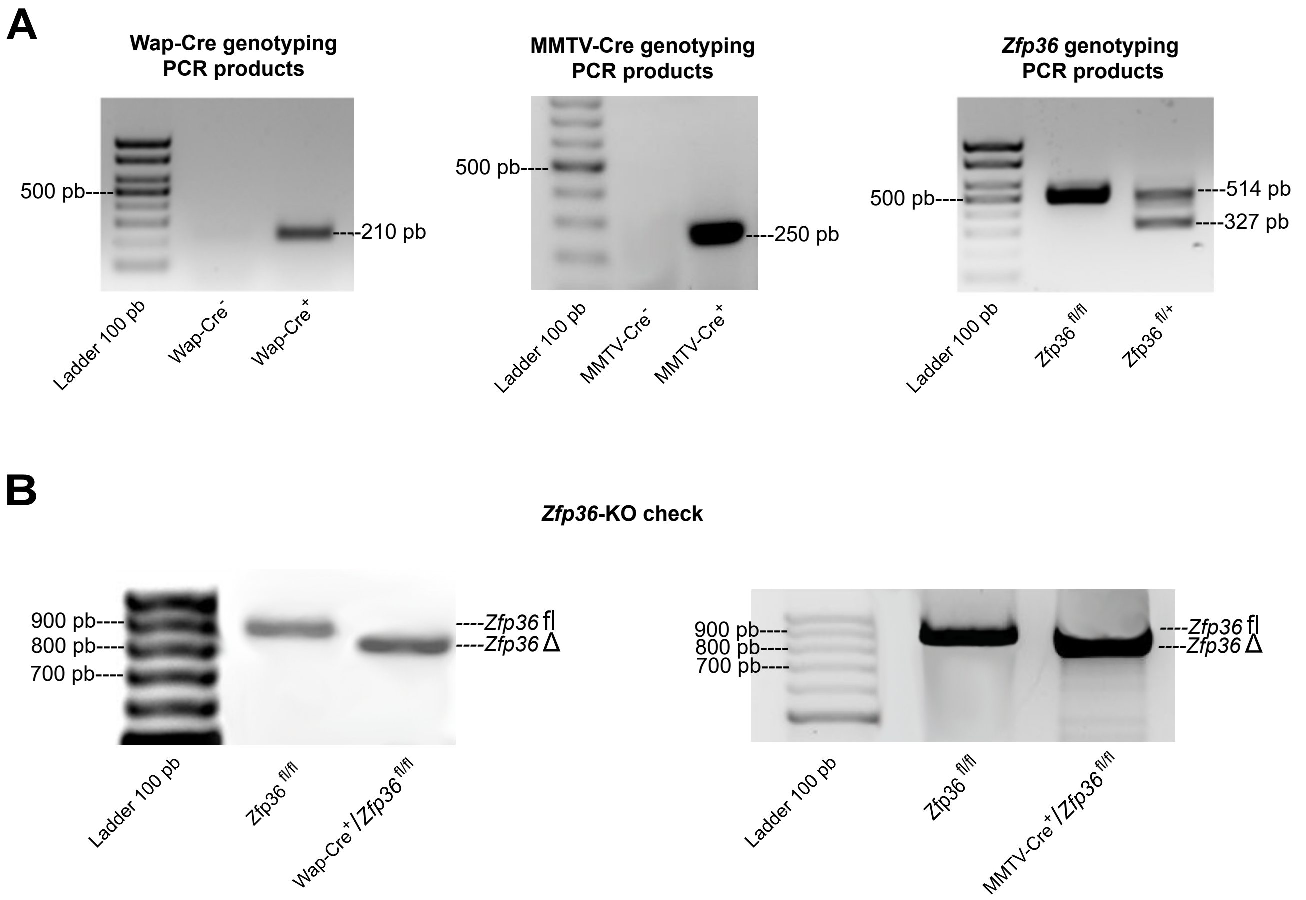
A
Wap-Cre genotyping PCR products
500 pb
210 pb
Ladder 100 pb
Wap-Cre-
Wap-Cre+
MMTV-Cre genotyping PCR products
500 pb
250 pb
Ladder 100 pb
MMTV-Cre-
MMTV-Cre+
Zfp36 genotyping PCR products
500 pb
514 pb
327 pb
Ladder 100 pb
Zfp36 fl/fl
Zfp36 fl/+
B
Zfp36-KO check
900 pb
800 pb
700 pb
Zfp36 fl
Zfp36 Δ
Ladder 100 pb
Zfp36 fl/fl
Wap-Cre+/Zfp36 fl/fl
900 pb
800 pb
700 pb
Zfp36 fl
Zfp36 Δ
Ladder 100 pb
Zfp36 fl/fl
MMTV-Cre+/Zfp36 fl/fl

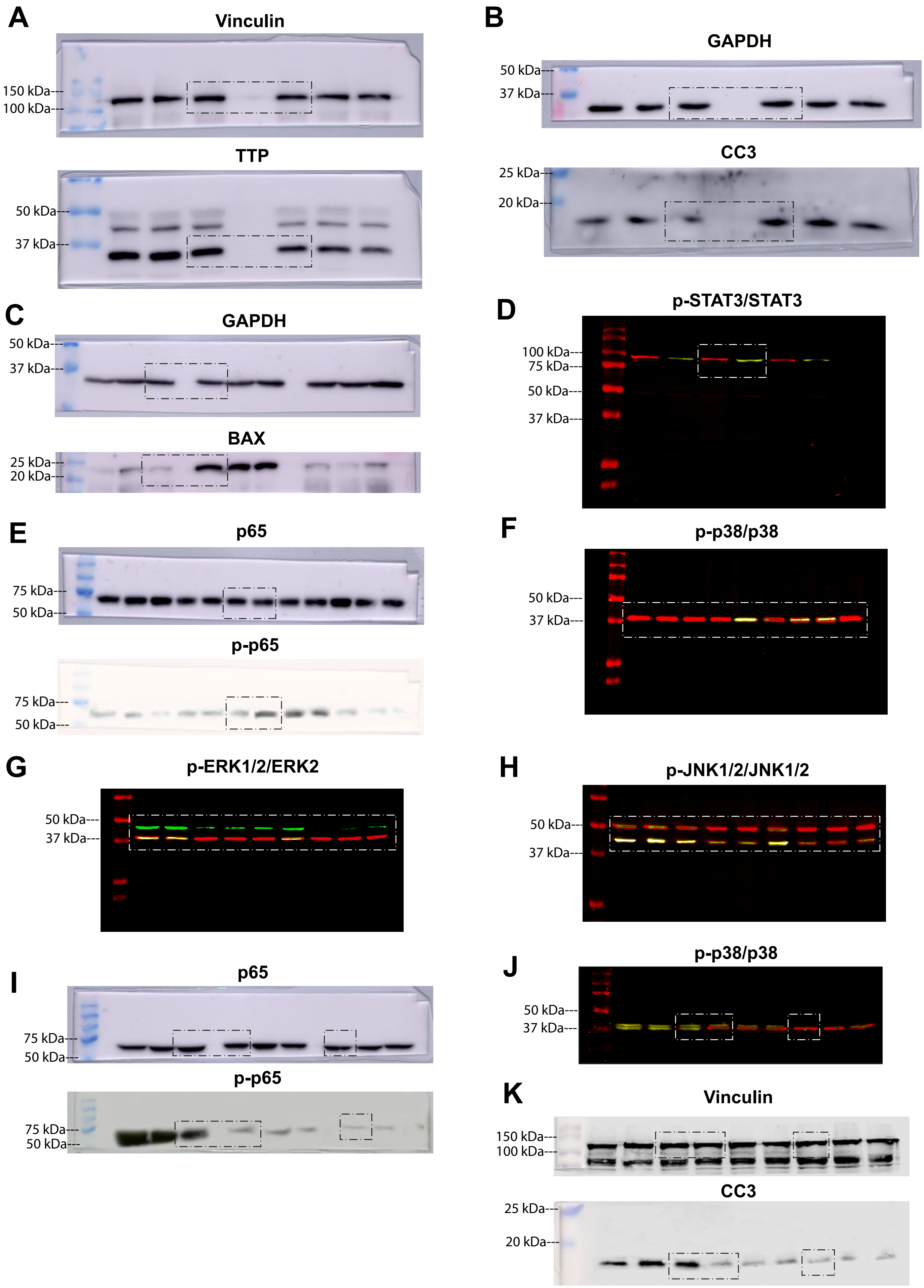

A
Vinculin
150 kDa
100 kDa
TTP
50 kDa
37 kDa
B
GAPDH
50 kDa
37 kDa
CC3
25 kDa
20 kDa
C
GAPDH
50 kDa
37 kDa
BAX
25 kDa
20 kDa
D
p-STAT3/STAT3
100 kDa
75 kDa
50 kDa
37 kDa
E
p65
75 kDa
50 kDa
p-p65
75 kDa
50 kDa
F
p-p38/p38
50 kDa
37 kDa
G
p-ERK1/2/ERK2
50 kDa
37 kDa
H
p-JNK1/2/JNK1/2
50 kDa
37 kDa
I
p65
75 kDa
50 kDa
p-p65
75 kDa
50 kDa
J
p-p38/p38
50 kDa
37 kDa
K
Vinculin
150 kDa
100 kDa
CC3
25 kDa
20 kDa

**SUPPLEMENTARY TABLES**

| Primer ID | Sequence (5´->3´) |
|---|---|
| TTP (g)_P1_Fw | GAACCCTCTCTCGATCGGGGATAC |
| TTP (g/KO) _P2_Rv | GGATGGAGTCCGAGTTTATGTTCCAA |
| TTP (KO) _P3_Fw | CTGGCTGGAAATGAGAGAGG |
| TTP (KO) _P4_Rv | CACCCCTTACGCCAGAACTA |
| WAP-CRE_W003 _Fw | TAGAGCTGTGCCAGCCTCTTC |
| CRE_C031_ RV | CATCACTCGTTGCATCGACC |
| MMTV-CRE_Fw | CTGATCTGAGCTCTGAGTG |

**Table 1:** Primer sequences used to genotype transgenic mice strains.

| Transgene | Product´s size (pb) | Primers used | Aim |
|---|---|---|---|
| WAP-Cre$^{+}$ | 210 | W003/C031 | Rutinary genotyping |
| MMTV-Cre$^{+}$ | 250 | MMTV-CRE/C031 | |
| *Zfp36*$^{+/+}$ | 327 | P1/P2 | |
| *Zfp36*$^{fl/+}$ | 514/327 | P1/P2 | |
| *Zfp36*$^{fl/fl}$ | 514 | P1/P2 | |
| *Zfp36*$^{+/+}$ | 683 | P3/P2 | *Zfp36* KO evaluation |
| *Zfp36*$^{fl/+}$ | 870/683 | P3/P2 | |
| *Zfp36*$^{fl/fl}$ | 870 | P3/P2 | |
| *Zfp36Δ* | 769 | P3/P4 | |

**Table 2:** Size of genotyping PCR products and combination of primers used.

| Primer ID | Sequence (5´-> 3´) | Melting temperature (°C) |
|---|---|---|
| *Zfp36 (r)_Fw* | CCACCTCCTCTCGATACA | 60 |
| *Zfp36 (r)_Rv* | GCTTGGCGAAGTTCACCCA | |
| *Tnfα_Fw* | CCACCACGCTCTTCTGTCTACT | 60 |
| *Tnfα_Rv* | GGGTCTGGGCCATAGAACTGAT | |
| *Il-6_Fw* | ATCCAGTTGCCTTCTTGGGA | 62 |
| *Il-6_Rv* | CCAGTTTGGTAGCATCCATCA | |
| *Lif_Fw* | CAGCGCCAATGCTCTCTTCATT | 62 |
| *Lif_Rv* | ATATTGGTCAGGGAGGCGCT | |

| | | |
|---|---|---|
| *Dusp6* | CTCGGATCACTGGAGCCAAA | 60 |
| *Dusp6* | GACAGAGCGGCTGATACCTG | 60 |
| *RNA18s_Fw* | GTAACCCGTTGAACCCCACCATT | 60 |
| *RNA18s_Fw* | CATCCAATCGGTAGTAGCG | |

**Table 3:** Primer´s sequences used in RT- qPCR analyses and melting temperatures employed.

| Antibody WB | Manufacturer | Catalogue number | Concentration | Species |
|---|---|---|---|---|
| TTP | Lab. Blacksear | ------ | 1:1000 | Rabbit |
| Vinculin | Santa Cruz | Sc-73614 | 1:5000 | Mouse |
| Cleaved caspase 3 | Cell Signaling | 9661 | 1:1000 | Rabbit |
| BAX | Santa Cruz | Sc-493 | 1:1000 | Rabbit |
| STAT3 total | Cell Signaling | 9139 | 1:1000 | Mouse |
| Fosfo-STAT3 (Tyr705) | Cell Signaling | 9145s | 1:1000 | Rabbit |
| GAPDH | Santa Cruz | Sc-322233 | 1:5000 | Mouse |
| p65 total | Cell Signaling | 8242 | 1:1000 | Rabbit |
| Fosfo-p65 (Ser536) | Cell Signaling | 3031 | 1:1000 | Rabbit |
| ERK2 total | Santa Cruz | Sc-1647 | 1:1000 | Mouse |
| Fosfo-ERK ½ (Tyr204) | Cell Signaling | 4370 | 1:1000 | Rabbit |
| JNK ½ total | Cell Signaling | 9258 | 1:1000 | Rabbit |
| Fosfo-JNK ½ (Thr183/Tyr185) | Cell Signaling | 9255 | 1:1000 | Mouse |
| p38 total | Cell Signaling | 92125 | 1:1000 | Rabbit |
| Fosfo-p38 (Thr180/Tyr 182) | Cell Signaling | 92165 | 1:1000 | Mouse |
| IκBα | Cell Signaling | 9242 | 1:1000 | Rabbit |
| Actina | Santa Cruz | Sc-1616R | 1:1000 | Rabbit |

**Table 4:** Antibodies used in WB analysis, their manufacturer, catalogue number, dilution employed and species in which they were made.

| Antibody | Manufacturer | Catalogue number | Dilution | Species |
|---|---|---|---|---|
| Cleaved caspase 3 | Cell Signaling | 9661 | 1:200 | Rabbit |
| Fosfo-p38 (Thr 180/ Tyr 182) | Cell Signaling | 4631 | 1:200 | Rabbit |
| Fosfo-p65 (Ser 536) | Cell Signaling | 3031s | 1:200 | Rabbit |
| Fosfo-STAT 3 (Tyr 705) | Cell Signaling | 9145s | 1:200 | Rabbit |
| TTP | Lab Balckshear | ------- | 1:200 | Rabbit |

**Table 5:** Antibodies used in IFs, their manufacturer, catalogue number, dilution employed and species in which they were made.
